# Multi-algorithm particle simulations with Spatiocyte

**DOI:** 10.1101/079293

**Authors:** Satya N. V. Arjunan, Koichi Takahashi

## Abstract

As quantitative biologists get more measurements of spatially regulated systems such as cell division and polarization, simulation of reaction and diffusion of proteins using the data is becoming increasingly relevant to uncover the mechanisms underlying the systems. Spatiocyte is a lattice-based stochastic particle simulator for biochemical reaction and diffusion processes. Simulations can be performed at single molecule and compartment spatial scales simultaneously. Molecules can diffuse and react in 1D (filament), 2D (membrane) and 3D (cytosol) compartments. The implications of crowded regions in the cell can be investigated because each diffusing molecule has spatial dimensions. Spatiocyte adopts multi-algorithm and multi-timescale frameworks to simulate models that simultaneously employ deterministic, stochastic and particle reaction-diffusion algorithms. Comparison of light microscopy images to simulation snapshots is supported by Spatiocyte microscopy visualization and molecule tagging features. Spatiocyte is open-source software and is freely available at http://spatiocyte.org.

## 1 Introduction

Heterogeneity and complex behavior observed at the cellular scale can arise from basic reaction-diffusion interactions at the molecular scale. Cell polarization, morphogenesis, chemotaxis and cytokinesis are some of the processes strongly coupled with the diffusion and spatiotemporal localization of signaling proteins. In addition to noisy reactions from low reactant numbers, the stochastic nature of the molecular interactions can also induce oscillations [1] and excite systems [2] in space and time. Consequently, spatial stochastic simulators have become important tools to elucidate the molecular mechanisms driving these processes [3–5]. Spatial simulators are also necessary for bottom-up construction of biophysically and biochemically realistic whole cell models [6]. Current simulators can be broadly categorized as (i) mesoscale, with the space discretized into subvolumes [7–12]; (ii) microscale, with each molecule tracked individually [13–19]; and (iii) hybridized meso-and microscale [20–22]. Mesoscopic simulators can advance time faster than microscale methods because they only track the concentration of species in the subvolumes instead of the exact position of each molecule. However, with the absence of such positional information, mesoscopic simulators cannot depict the effects of individual molecular interactions such as molecular crowding [17,19] and rebinding [15].

Spatiocyte is a hybrid macro-and microscale simulator for stochastic reaction-diffusion processes. Molecule species homogeneously distributed (HD) in a compartment are simulated at the macroscopic scale whereas heterogeneously distributed (non-HD) molecules are individually diffused in the lattice. Species found in many copies, which are usually evenly distributed in a compartment, can be simulated rapidly at the macroscopic scale by only tracking their molecule number without explicit diffusion steps. Spatiocyte can simulate reaction-diffusion of molecules on filaments such as microtubules, in addition to membranes and in solutions [23]. The simulator includes a visualizer that can display the position of non-HD molecules and their time-averaged trajectory, as visualized using a light microscope, for rapid visual comparison to experimentally captured images. Compartment geometry can be specified using a combination of geometric primitives such as cuboids, ellipsoids and cylinders, which can be translated and rotated. Spatiocyte can virtually tag and track a subpopulation of molecules individually, even as they transition from one state to another or between compartments. With this feature, it is possible to compare the diffusion behavior of the molecules with sparsely tagged molecules in experiments. Metabolic reactions usually involve a large number of molecules, most of which can be assumed to be HD. These reactions can be executed using Michaelis-Menten, Gillespie or mass action algorithms alongside Spatiocyte reaction-diffusion processes.

A detailed description of the Spatiocyte particle simulation algorithm is provided by Arjunan and Tomita [24]. Briefly, the space is discretized into hexagonal close-packed (HCP) lattice with regular sphere voxels. Each voxel can be occupied by a single non-HD molecule as the voxel size approximates the size of the molecule. A diffusing non-HD molecule can walk to one of its 12 neighbor voxels by random selection in a diffusion step interval. The interval is calculated from the species diffusion coefficient and the voxel radius. The walk is successful if the target voxel is vacant; otherwise, if it contains a reactant pair, a collision occurs and the molecules react with a probability corresponding to the reaction rate constant. If the voxel is occupied by a non-reactive molecule, the source molecule stays in the original voxel. HD molecules react with non-HD molecules in an event-driven manner according to a spatially adapted Next Reaction method [24–25]. The diffusion and reaction processes of the simulation method have been verified with analytical and numerical solutions [24].

Using Spatiocyte and protein measurements such as concentration, binding, dissociation and diffusion constants in different states (monomer, homodimer, heterodimer, cytosolic, membrane-bound, etc.), we can build a quantitative model that closely represents the cellular system of interest. We can verify the model by comparing simulation outcomes with wild type and mutant phenotypes of the system. Often, not all model parameters can be measured and in such cases, the parameters can be estimated by adjusting them until the simulated phenotypes agree with observations. Since Spatiocyte simulations are rapid, we can use the model to explore the parameter space to predict the system behavior with different combinations of parameter values. For example, we can predict the phenotype with varying expression and activity levels of proteins (e.g., phosphorylation and dephosphorylation), protein mutation (removal of one or more of its functions), cell morphology and initial conditions. These predictions can be tested experimentally. If what we observe in experiments do not match the predictions, the model can be adjusted until it recapitulates the observations. The adjustments will provide new insights about the molecular mechanisms underlying the system. In a more advanced case, Spatiocyte can also be used to design models, made up of components with well characterized parameters, to generate a particular behavior of a system (e.g., spatial pattern or domain formation). The model can then be realized experimentally to generate such behaviors.

## 2 Materials

### 2.1 Input model parameters

Spatiocyte requires simulation model parameters to write the model and perform simulations. The model parameters include the simulation voxel size, compartment geometry and dimensions, molecule species, species molecule number and diffusion coefficient, reactions and their rate constants. In the Methods section, the steps to build a multi-algorithm particle simulation model are provided. There are also several other example models included in the software package.

### 2.2 Spatiocyte software

Spatiocyte runs stably on Ubuntu Linux but it is still experimental on Mac OS X and Windows systems. Up-to-date download and installation instructions of Spatiocyte can be found at http://spatiocyte.org. On a fresh Ubuntu system, Spatiocyte requires additional libraries and packages to run. These include Python, Boost.Python, Git, Hierarchical Data Format 5, Matplotlib, NumPy, SciPy and GNU Scientific Library (GSL), all of which are automatically installed along with Spatiocyte. After the model is parsed, the simulator creates the compartments according to the specified geometry and populates the molecules as given in the initial conditions. Logger modules are also initialized and they start logging simulation data in the specified log files. After that, graphical or command line interface is activated to transfer the control of program execution to the user. The user can run the simulation for a given time and view the dynamics with the Spatiocyte Visualizer. The visualizer is a separate program in the Spatiocyte package that can run concurrently with the simulator. It loads the log file created by the visualization logger modules to display time-lapse molecule positions or simulated microscopy snapshots using the OpenGL library. Screenshots of the simulation can be saved in Portable Network Graphics (PNG) format and animated to create high resolution movies.

## 3 Methods

### 3.1 Build a simulation model

Here, we will build a multi-algorithm simulation model in Python. The model can also be written C++ but for simplicity, we focus on Python (*see* **Note 1**). The algorithm modules, such as diffusion, reaction, logger, tagger and molecule population can be specified as necessary in a model. Each module has its own set of options that are defined in the model by the user. These options are described in the general guide to build Spatiocyte models [26]. Our multi-algorithm model consists of mass action, Spatiocyte next-reaction and lattice-based particle reaction-diffusion methods.

1. *Prepare model parameters*. The parameters of the model are listed in Fig. 1. In the model, we have non-HD species, A, B and C, with molecules individually diffused, and HD species, E, S, ES, and P. We use mass action to simulate three reversible Michaelis-Menten type reactions, where a product, P is formed by a single enzyme-substrate complex, ES from a single substrate, S and an enzyme, E: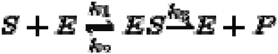. The product P will stochastically bind A to generate a heterodimer B with Spatiocyte next-reaction method: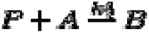. The fifth and final reaction, which is diffusion-influenced, involves reactant B that will bind another diffusing reactant A to generate C: 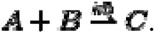
2. *Open an empty model file*. To write the model, open an editor and save an empty file as ode-snrp-particle.py. Add the Python code as provided in the steps below to the file and save as we go along. The complete model file is included in the examples/ode-snrp-particle directory of the Spatiocyte package.
3. *Specify steppers and voxel radius*. In the model, we refer to the Spatiocyte simulator as `theSimulator`. Since the term is used frequently throughout the code, we define sim as a short alias for theSimulator below. All Spatiocyte models require the SpatiocyteStepper. It advances the time steps of logger, reaction and diffusion modules in an event-driven manner. We set the radius of the hexagonal close-packed lattice voxels with the VoxelRadius option (*see* **Note 2**). Below, we have set the radius to 4.4 nm. Since we are building a multi-algorithm model consisting of mass action reactions, we also need to specify a stepper for ordinary differential equation solver called ODEStepper to execute the reactions in continuous-time. The remaining two reactions, next-reaction and diffusion-influenced reaction, are performed in discrete-time by the SpatiocyteStepper. We set the maximum step interval of the ODEStepper to 0.001 s for accuracy (*see* **Note 3**). Each compartment in the Spatiocyte model must have its StepperID assigned to the SpatiocyteStepper ID. This model only consists of the root compartment, rootSystem and we set its StepperID to SpatiocyteStepper id, 'SS' (*see* **Note 4**).

~~~
sim = theSimulator
s = sim.createStepper('SpatiocyteStepper', 'SS')
s.VoxelRadius = 4.4e-9
s = sim.createStepper('ODEStepper', 'DE')
s.MaxStepInterval = 1e-3
sim.rootSystem.StepperID = 'SS'
~~~
4. *Set compartment geometry*. We can specify the geometry of the root compartment by setting the GEOMETRY variable to one of the six supported geometric primitives: cuboid (‘0’), ellipsoid (‘1’), cylinder (‘2’), rod (‘3’), pyramid (‘4’) and erythrocyte (‘5’). For simplicity, we use a cuboid compartment geometry in this model. More complicated geometries can be constructed using a combination of the existing primitives. The three variables LENGTH[X, Y, Z] specify the compartment lengths in the direction of [x, y, z]-axis, respectively. Here, the lengths are set to 500 nm.

~~~
sim.createEntity('Variable', 'Variable:/:GEOMETRY').Value = 0
sim.createEntity('Variable', 'Variable:/:LENGTHX').Value = 5e-7
sim.createEntity('Variable', 'Variable:/:LENGTHY').Value = 5e-7
sim.createEntity('Variable', 'Variable:/:LENGTHZ').Value = 5e-7
~~~
5. *Define species, type and initial molecule numbers*. Each compartment is initially made up of empty voxels. We sometimes need to refer to these empty voxels in the simulation (*see* **Note 5**). These empty voxels are called VACANT species in the model and each compartment must have one such species declared to represent its empty voxels. By default, all species are non-HD type unless we set the Name property of the species to HD, as we have done for E, S, ES and P below. We have also set the species A, E and S to have 1500, 100 and 1000 molecules initially.

~~~
sim.createEntity('Variable', 'Variable:/:VACANT')
sim.createEntity('Variable', 'Variable:/:A').Value = 1500
sim.createEntity('Variable', 'Variable:/:B').Value = 0
sim.createEntity('Variable', 'Variable:/:C').Value = 0
v = sim.createEntity('Variable', 'Variable:/:E')
v.Value = 100
v.Name = 'HD'
v = sim.createEntity('Variable', 'Variable:/:S')
v.Value = 1000
v.Name = 'HD'
v = sim.createEntity('Variable', 'Variable:/:ES')
v.Value = 0 v.Name = 'HD'
v = sim.createEntity('Variable', 'Variable:/:P')
v.Value = 0
v.Name = 'HD'
~~~
6. *Populate non-HD species initialized with non-zero molecules*. In the model, only A is a non-HD species that has a non-zero initial number of molecules. We need to specify how to populate these explicitly represented molecules in the compartment using MoleculePopulateProcess. By default, the process will populate the initial 1500 molecules of A randomly in the compartment (*see* **Note 6** for alternative ways to populate). In the first line below, we have created a MoleculePopulateProcess object called ‘Process:/:pop’. In the second line, we have connected the species A to the process by adding the reference of the variable to the process’ VariableReferenceList. The first field (denoted here as ‘_’) specifies a name of the variable reference that will be used to identify it locally in the process. MoleculePopulateProcess does not have any predefined variable reference name to identify connected variables, so we have just given an empty name field, ‘_’. The second field specifies the path, ‘:/:’ and identity, A of the variable, which we have written here as ‘Variable:/:A’. More details on how to connect variables to processes are provided in the E-Cell System manual available at https://ecell3.readthedocs.io/en/latest/modeling.html.

~~~
p = sim.createEntity('MoleculePopulateProcess', 'Process:/:pop')
p.VariableReferenceList = [['_', 'Variable:/:A']]
~~~
7. *Set diffusion coefficient of non-HD species*. The DiffusionProcess is the module that specifies the diffusion properties of a species. We diffuse all three non-HD species, A, B and C in the compartment with a diffusion coefficient of 0.0005 μm^2^s^−1^.

~~~
d = sim.createEntity('DiffusionProcess', 'Process:/:d1')
d.VariableReferenceList = [['_', 'Variable:/:A']]
d.D = 5e-16
d = sim.createEntity('DiffusionProcess', 'Process:/:d2')
d.VariableReferenceList = [['_', 'Variable:/:B']]
d.D = 5e-16
d = sim.createEntity('DiffusionProcess', 'Process:/:d3') d.VariableReferenceList = [['_', 'Variable:/:C']]
d.D = 5e-16
~~~
8. *Define the reactions*. The three deterministic reactions involving HD species are performed by MassActionProcess. We assign the ODEStepper to the reaction module by setting the StepperID to DE. SpatiocyteNextReactionProcess executes the stochastic reaction that generates B when P and A react. DiffusionInfluencedReactionProcess performs the bimolecular reaction between the two diffusing non-HD species, A and B to produce C (*see* **Note 7**). For the first mass action reaction E + S→ES, we need to connect the species E, S and ES to a MassActionProcess object. As described in Step 6, we connect them by adding the references of the variables into the process’ VariableReferenceList. Note that each new line of the VariableReferenceList with the operator ‘=’ below does not overwrite the reference given in the previous line but adds the new reference to the existing list. Unlike in Step 6, in the second field of the variable reference we have used the relative path (w.r.t. the compartment path) of the variable, ‘:.:’ instead of the absolute path, ‘:/:’. Relative path is useful when we want to skip updating the paths of the variables when we change the name of the compartment. The third field denotes whether the variable is a substrate (‘-1’) or a product (‘1’) of the reaction. The remaining reactions follow the same conventions to connect the species to the corresponding processes.

~~~
# E + S --> ES
r = sim.createEntity('MassActionProcess', 'Process:/:r1') r.StepperID = 'DE'
r.VariableReferenceList = [['_', 'Variable:.:E','-1']]
r.VariableReferenceList = [['_', 'Variable:.:S','-1']]
r.VariableReferenceList = [['_', 'Variable:.:ES','1']]
r.k = 1e-22
# ES --> E + S
r = sim.createEntity('MassActionProcess', 'Process:/:r2')
r.StepperID = 'DE'
r.VariableReferenceList = [['_', 'Variable:.:ES', '-1']]
r.VariableReferenceList = [['_', 'Variable:.:E', '1']]
r.VariableReferenceList = [['_', 'Variable:.:S', '1']]
r.k = 1e-1
# ES --> E + P
r = sim.createEntity('MassActionProcess', 'Process:/:r3')
r.StepperID = 'DE'
r.VariableReferenceList = [['_', 'Variable:.:ES', '-1']]
r.VariableReferenceList = [['_', 'Variable:.:E', '1']]
r.VariableReferenceList = [['_', 'Variable:.:P', '1']]
r.k = 1e-1
# P + A --> B
r = sim.createEntity('SpatiocyteNextReactionProcess', 'Process:/:r4')
r.VariableReferenceList = [['_', 'Variable:/:P', '-1']]
r.VariableReferenceList = [['_', 'Variable:/:A', '-1']]
r.VariableReferenceList = [['_', 'Variable:/:B', '1']]
r.k = 5e-24
# A + B --> C
r = sim.createEntity('DiffusionInfluencedReactionProcess', 'Process:/:r5')
r.VariableReferenceList = [['_', 'Variable:/:A', '-1']]
r.VariableReferenceList = [['_', 'Variable:/:B', '-1']]
r.VariableReferenceList = [['_', 'Variable:/:C', '1']]
r.k = 5e-24
~~~
9. *Specify data loggers and the simulation time*. Below we use VisualizationLogProcess to log the coordinates of A, B and C in lattice every 0.1 s in a binary format log file called VisualLog.dat (*see* **Note 8**). The Spatiocyte Visualizer can load the log file to display the 3D position of the molecules in time while the simulation is running or after it has ended. We set the IteratingLogProcess to record each species’ molecule number every 0.01 s from the beginning of the simulation until 99 s in a csv format file, IterateLog.csv (*see* **Note 9**). Finally, we can tell the simulator how long to run the model. Here, we set it to run for 100 s.

~~~
l = sim.createEntity('VisualizationLogProcess', 'Process:/:l1')
l.VariableReferenceList = [['_', 'Variable:/:A']]
l.VariableReferenceList = [['_', 'Variable:/:B']]
l.VariableReferenceList = [['_', 'Variable:/:C']]
l.LogInterval = 1e-1
l = sim.createEntity('IteratingLogProcess', 'Process:/:l2')
l.VariableReferenceList = [['_', 'Variable:/:A']]
l.VariableReferenceList = [['_', 'Variable:/:B']]
l.VariableReferenceList = [['_', 'Variable:/:C']]
l.VariableReferenceList = [['_', 'Variable:.:E']]
l.VariableReferenceList = [['_', 'Variable:.:S']]
l.VariableReferenceList = [['_', 'Variable:.:ES']]
l.VariableReferenceList = [['_', 'Variable:.:P']]
l.LogInterval = 1e-2
l.LogEnd = 99
run(100)
~~~

**Fig 1.**
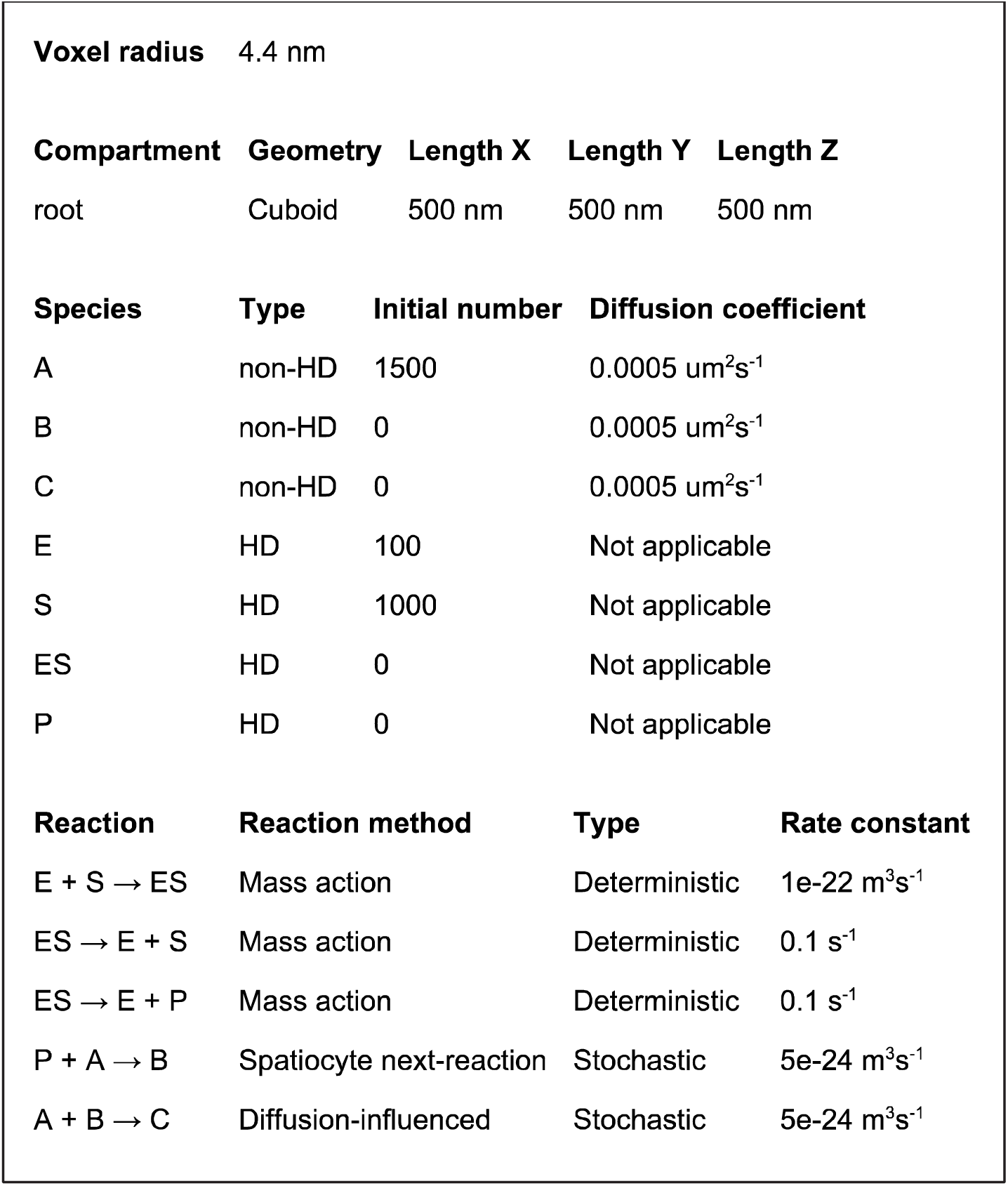
Input parameters of the multi-algorithm simulation model.

### 3.2 **Run the model**

After successfully installing Spatiocyte (*see* **Note 10**), we can simulate the multi-algorithm model in a terminal by issuing:

~~~
$ ecell3-session ode-snrp-particle.py
~~~

The simulator will run for 100 s and terminate. In the current working directory, it would have saved two log files, VisualLog.dat and IterateLog.csv.

### 3.3 **Display simulation results**

Finally, we can visualize the data logged by the two loggers with the following steps.

1. *View diffusing molecules*. Even while the simulation is running we can view the dynamics of the diffusing molecules with Spatiocyte Visualizer (Fig. 2) by issuing

~~~
$ spatiocyte VisualLog.dat
~~~

in the working directory. The visualizer will load VisualLog.dat and display the molecule positions of non-HD species, A, B and C as displayed in Fig. 3. The shortcut keys to control the visualizer are provided in the Spatiocyte guide [26]. For example, the right arrow key will advance the time forward whereas the left arrow key, backward. Pressing the space bar key will pause or resume the time advancement.
2. *View time course profiles*. From IterateLog.csv, we can plot the time course profiles of the logged species using a helper Python script called plotIterateLog.py, which is included in the Spatiocyte examples/plot directory. Copying the file into the working directory and issuing the command below will display the profiles as shown in Fig. 4.

~~~
$ python plotIterateLog.py
~~~

**Fig 2.**
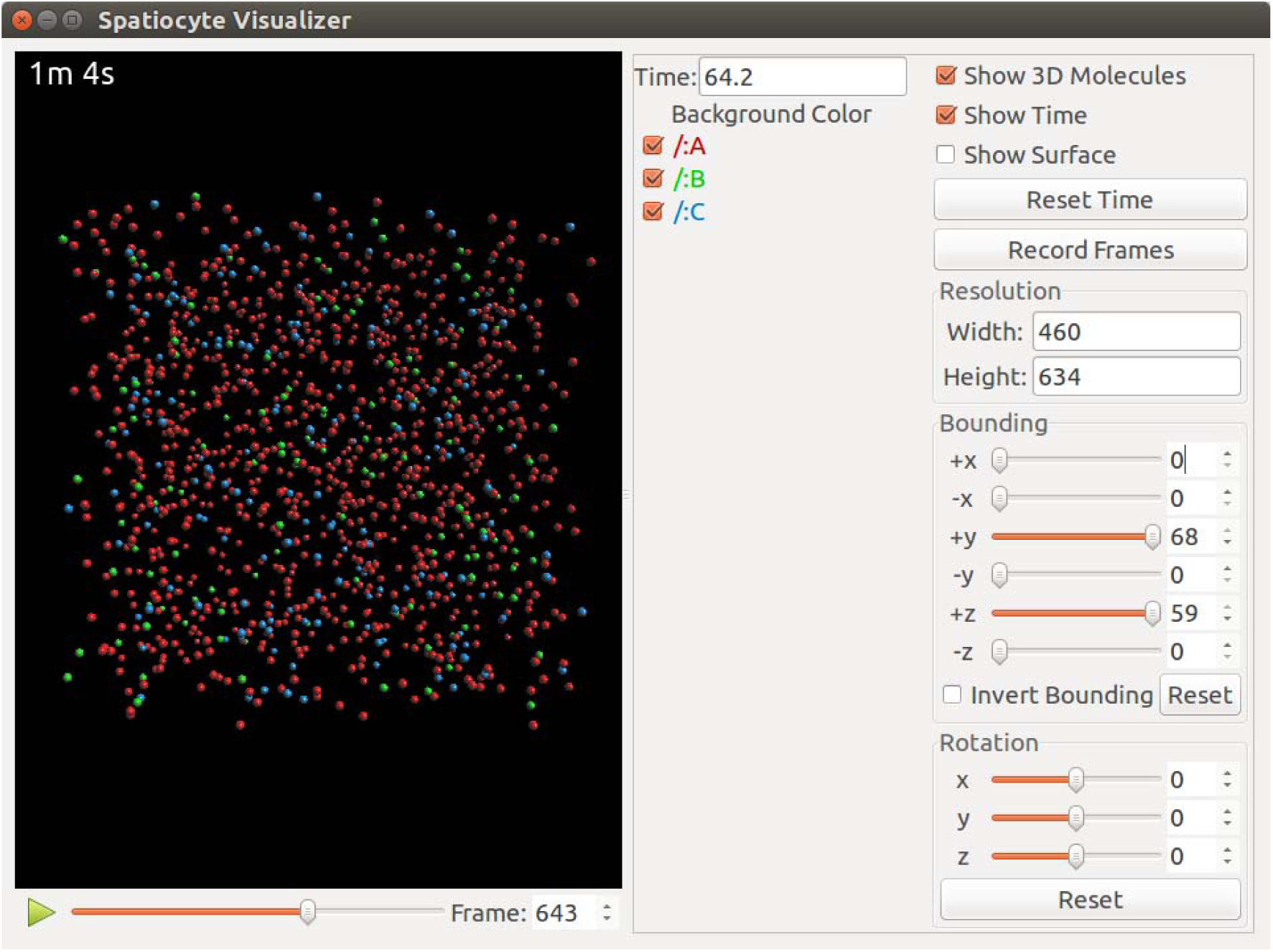
The graphical user interface of Spatiocyte Visualizer

**Fig 3.**
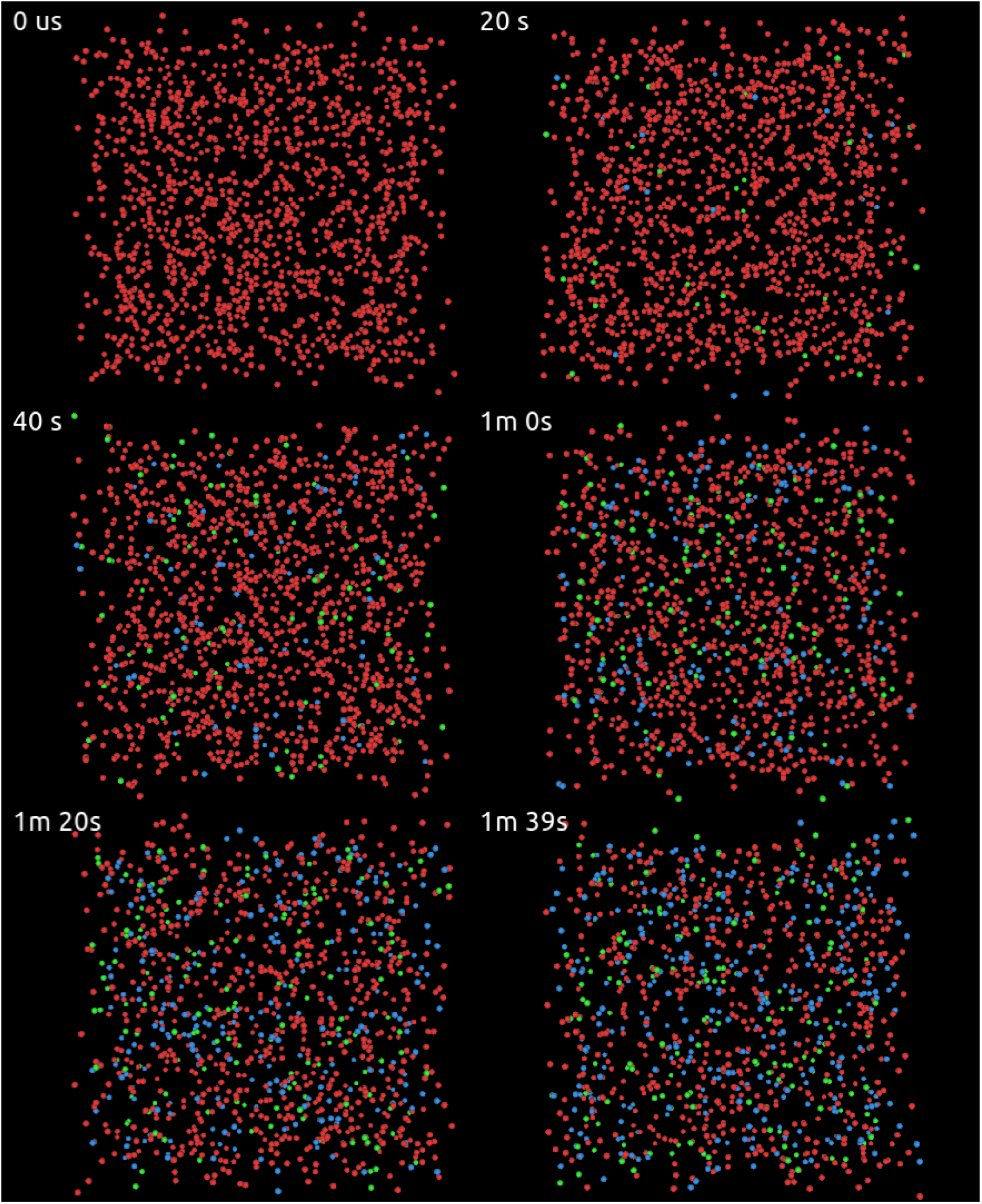
Simulation snapshots of the multi-algorithm model. Initially, all 1500 molecules of A (red) are populated randomly in cubic space. As time advances, more B (green) and C (blue) molecules start to appear while A decreases as a result of the multi-algorithm reactions.

**Fig 4.**
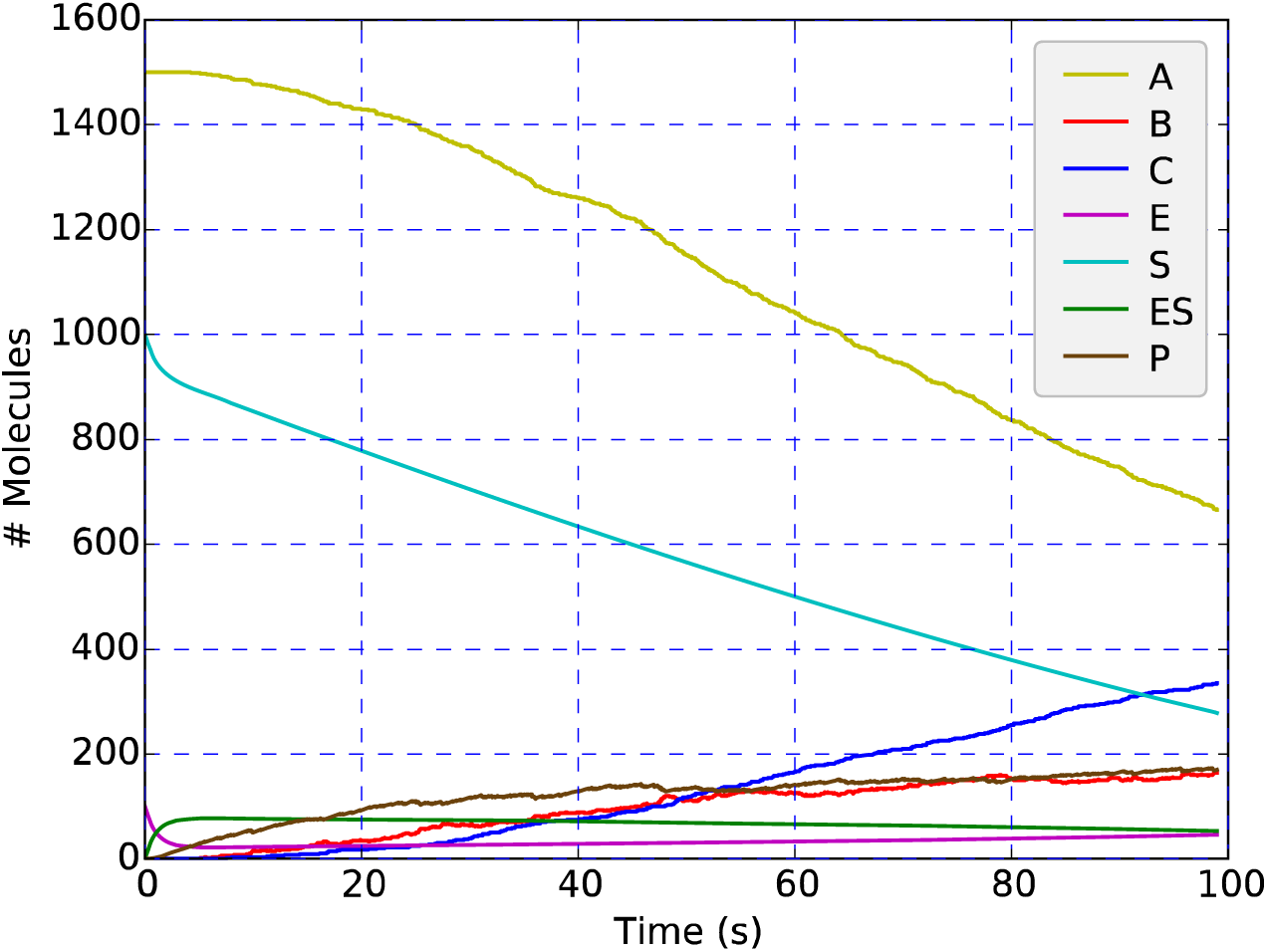
Time course profiles of the multi-algorithm simulation. S, E and ES show smooth lines over time because they are only involved in deterministic reactions. P which is involved in both stochastic and deterministic reactions, displays noisy increase over time.

## 4 Case Studies

We have previously used Spatiocyte to model *Escherichia coli* division site regulators, MinD and MinE proteins that periodically cycle the poles of the rod-shaped bacterium [24]. Our model is the first to corroborate the prediction that MinE can bind to the membrane independently using its membrane domain [27–28] after it is recruited from the cytoplasm by MinD. The model also first predicted that independently membrane-bound MinE can rebind with other MinD’s on the membrane. These predictions were later supported experimentally [29–30]. Recently, we built a multi-algorithm simulation model of erythrocyte band 3 membrane cluster formation with Spatiocyte [31]. The model showed that strong affinity between the clustering molecules and irreversibly binding hemichromes aid the generation of oxidation induced clusters as observed in experiments. The simulated cluster size increased towards an irreversible state when oxidative stress is introduced repeatedly. The model also predicted that erythrocytes with deficient spectrin cytoskeletal filaments have more and larger band 3 clusters. In addition, together with our colleagues, we have recently developed a bioimaging simulation framework that produces simulated microscopy images from 3D molecule coordinates generated by particle simulators such as Spatiocyte [32]. The simulated images can be compared with actual microscopy images at the level of photon-counting units. We verified the bioimaging simulator by comparing simulated images of several in vitro and in vivo Spatiocyte models with experimentally obtained microscopy images.

Here, as another example of Spatiocyte application, we show that *E. coli* cell geometry can regulate MinD oscillation period, while the cell size controls the peak MinD concentration on the membrane. Previous works have shown that MinD dynamics can be regulated by the geometry [33–37] and topology [38] of the membrane. Varma and colleagues [33] used E. coli lacking penicillin binding proteins to produce branched cells with three poles (Y-shaped) to investigate the effects of the mutant cell geometry on MinD membrane dynamics. Cells having almost equal branch lengths displayed non-reversing clockwise or counterclockwise rotational MinD polar localization. In cells where two of its poles are closer to each other than the third pole, MinD cycled back and forth symmetrically between the two poles and the third pole. Adapting our previously reported model [24], we investigated MinD dynamics in different geometric configurations of the branched cells as illustrated in Fig. 5, with fixed protein concentrations.**Fig. 6** displays the corresponding kymographs of MinD simulation results.

**Fig. 5.**
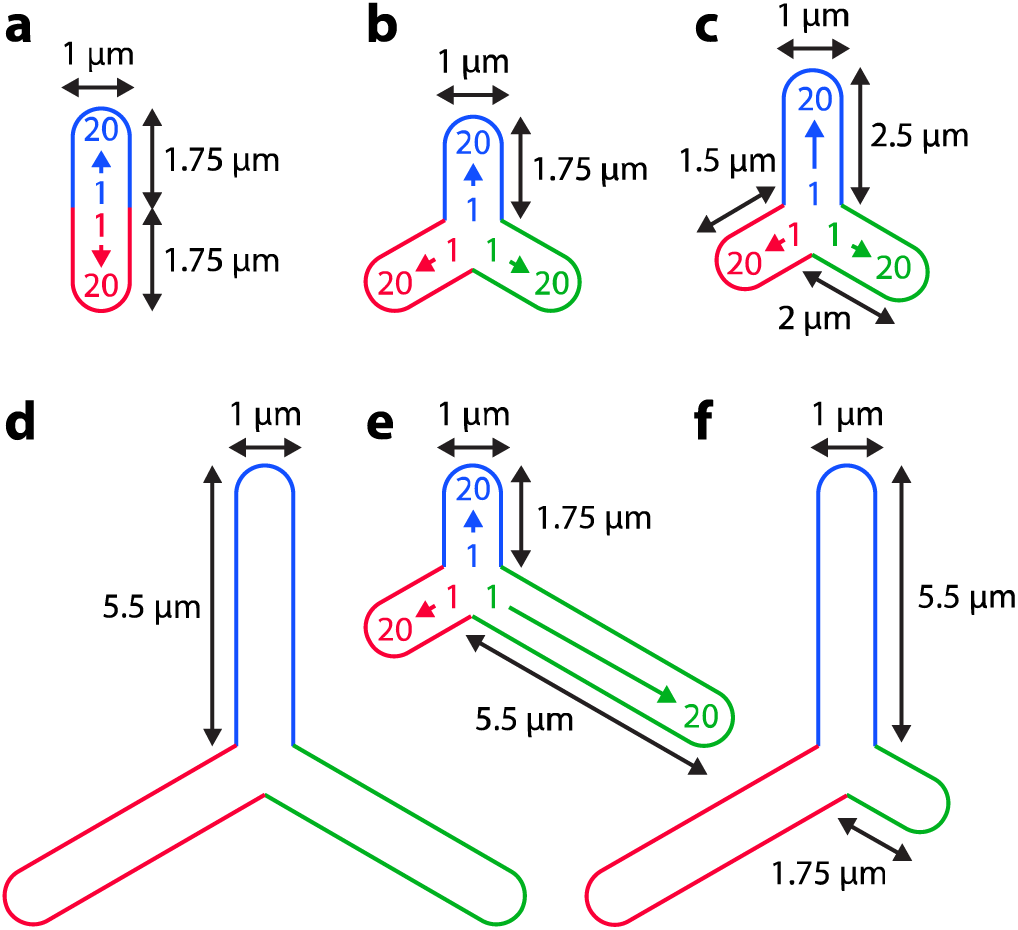
Schematic representation of *E. coli* geometric configurations used in simulations. Blue, red and green borders indicate the different branches of the cell. The range 1 to 20 in each branch represents the bin # in which the membrane concentration of MinD is calculated. The length of a branch is not stated if the length is already specified for another branch of the cell having the same length.

**Fig. 6.**
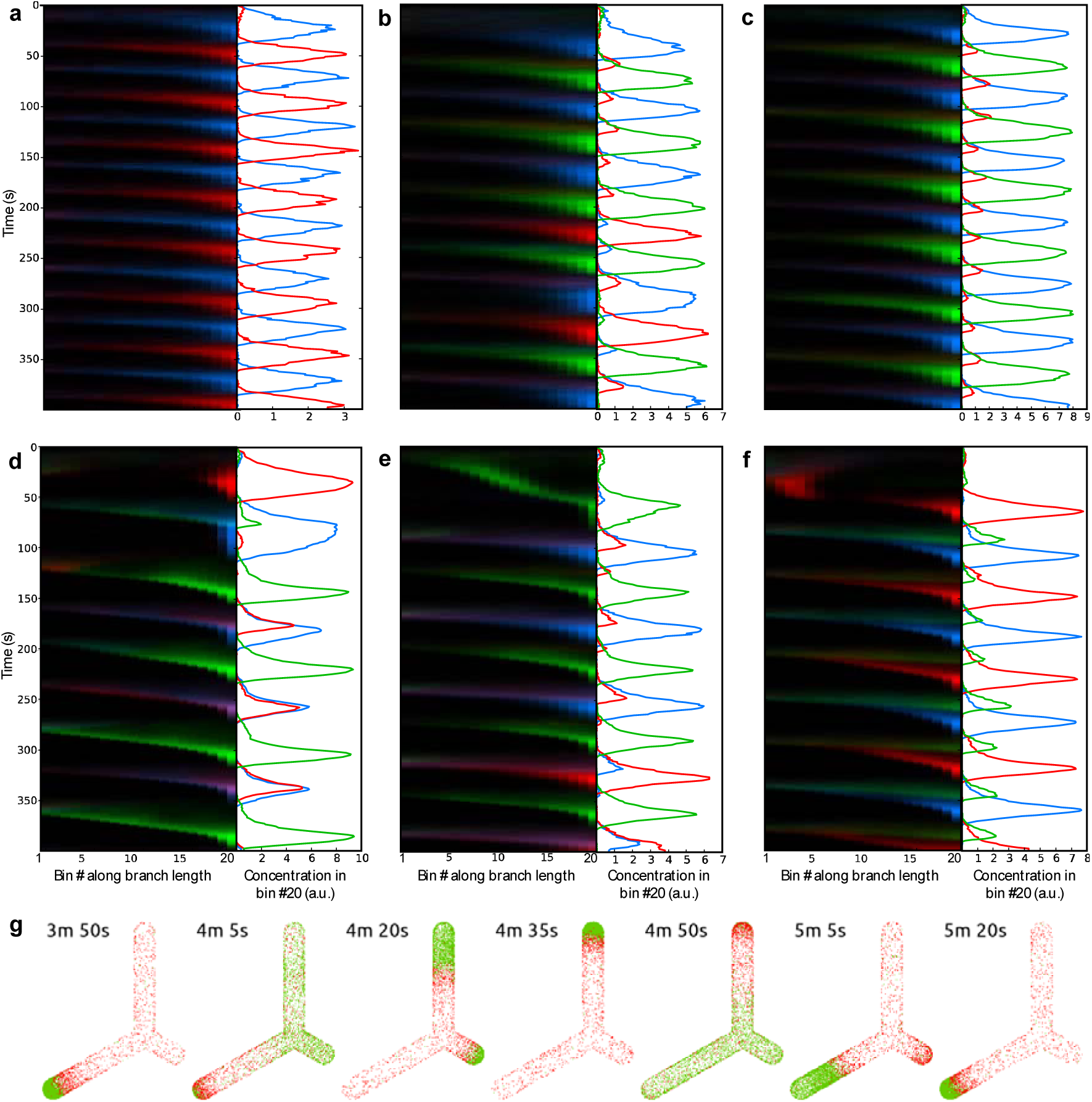
Simulation results of E. coli with different geometric configurations. (**a**)-(**f**) left panel: Kymograph of MinD concentration in cells with geometries specified in Fig. 5 (**a**)-(**f**), respectively; (**a**)-(**f**) right panel: MinD concentration in bin #20. Blue, red and green indicate MinD concentration corresponding to the branch color specified in Fig. 5; (**g**): Example simulation snapshots of MinD (green) and MinE (red) in cell (**f**), generated by Spatiocyte Visualizer.

In cells with equal branch lengths of 1.75 μm, MinD showed symmetrical oscillation that occasionally switched poles randomly. In all other configurations, MinD produced stable symmetrical oscillations. Despite implementing such diverse geometric configurations of the branched cells, our model is unable to recapitulate the rotational oscillation as observed in the experiments. Further detailed simulations are necessary to identify the requirements of rotational MinD oscillation in branched cells. Nonetheless, our preliminary simulations indicate that the period of oscillation increases as the total length of the branches increases. Regardless of the cell geometries, the peak concentration of MinD on the membrane correlates with the total surface area or the volume of the cell. More simulations and analyses are required to reveal how the oscillation period and MinD membrane concentration are regulated by the branch lengths, cell volume and membrane surface area.

## 5 Notes

1. *Modeling language*. Spatiocyte models can be written either in Python or C++. With Python, we can simulate the model without compiling it into an executable, which would take up additional time and effort. It is also easier with Python to perform multiple iterations of a model and introduce conditions when running the simulation because it is a scripting language. C++ models on other hand permit more flexibility and are useful when we want to optimize the compiled executable for a specific CPU for faster run times.
2. *VoxelRadius value*. For better simulation accuracy, the value of VoxelRadius should be close to the hydrodynamic radius of the diffusing species [24]. However, the simulation would consume more computation time when the VoxelRadius is small because of the shorter simulation time steps required when performing smaller diffusion steps over the voxels. The memory usage also increases linearly with the number of voxels. In a 64-bit system, each voxel typically takes up 108 bytes of memory. The number of voxels with radius, *r* in a volume, *V* is given by *V*/(4*r*^3^2^0.5^). Therefore, in the initial stages of modeling, we usually first perform quick simulations with larger voxels and attempt to recapitulate experimentally observed phenotypes by modifying reactions and other unknown model parameter values. As the simulation phenotypes start to agree with observations, we gradually reduce the size of the voxels to the hydrodynamic radius of diffusing species.
3. *MaxStepInterval of ODEStepper*. ODEStepper executes mass action reactions at varying step intervals. To allow fast simulations, it dynamically increases the step interval when accuracy would not be compromised for the reactions. However, SpatiocyteStepper typically performs diffusion-influenced reactions and next-reaction at very short intervals because of the short diffusion time steps. To ensure that the molecule number of the species in the mass action reactions are valid at these short intervals when they are accessed by SpatiocyteStepper reactions, we set the MaxStepInterval of the ODEStepper to a small value.
4. *StepperID inheritance*. The StepperID for all modules in a compartment such as DiffusionProcess, MassActionProcess and SpatiocyteNextReactionProcess is inherited from the compartment’s StepperID. Since all modules except MassActionProcess are executed in event-driven manner by SpatiocyteStepper, we set the root compartment's StepperID to the SpatiocyteStepper ID. In each MassActionProcess module we can directly set its StepperID to the ODEStepper.
5. *Reactions involving VACANT species*. In some reactions, we need to specify the VACANT species of the compartment as a reactant. For example, in the diffusion-influenced membrane association reaction, where a cytosolic A binds to the membrane to form Am, the VACANT voxels of the membrane compartment is one of the reactants of the second-order reaction: A + membrane:VACANT --> Am.
6. *Populating molecules in a compartment*. Non-HD molecules are by default randomly populated throughout the compartment of the species with uniform distribution by MoleculePopulateProcess. We can also set a specific range to populate along each dimension of the compartment by setting the Origin[X,Y,Z] and Uniform[Length, Width, Height] options of the process. Molecules can also be populated along the length of the compartment divided into a given number of bins with different occupancy fractions using the LengthBinFractions array option. It specifies the number of bins and the population fraction of molecules over the total available vacant voxels in each bin.
7. *Reaction module selection*. We use DiffusionInfluencedReactionProcess only for second-order reactions where both reactants are diffusing non-HD species. If all the reactants of first-or second-order reaction are HD, we can use MassActionProcess. For all first-order reactions we can also use SpatiocyteNextReactionProcess. We implement SpatiocyteNextReactionProcess for all second-order reactions when either (or both) of the reactants is HD.
8. *VisualizationLogProcess default log interval*. If we do not specify the LogInterval value of VisualizationLogProcess, the logger will log the coordinates of its listed diffusing species at all SpatiocyteStepper time steps. This option is useful when we want to detect the exact time when a molecule changes its state in space.
9. *Log molecule coordinates in csv format*. Spatiocyte also comes with another logger module called CoordinateLogProcess that saves the coordinates of non-HD molecules at defined intervals in csv format. The coordinate data is useful for the user to perform custom detailed analysis of the simulation. The log file can also be read by a helper Python plotting script called plotCoordinateLog.py, included in the Spatiocyte examples directory.
10. *Verifying Spatiocyte installation*. To test if the Spatiocyte installation is successful, issue the following command in a terminal: $ ecell3-session The above command will start the Python command line interface of Spatiocyte. If for some reason, the interface does not come up, the error message can be posted to the Spatiocyte Users forum at https://groups.google.com/forum/?hl=en#!forum/spatiocyte-users for help.

## Acknowledgment

We thank Masaki Watabe, Hanae Shimo and Kaizu Kazunari for discussions that led to the improvement of Spatiocyte usage. We also appreciate Kozo Nishida for Spatiocyte software packaging, installation and documentation assistance.

